# Saturation mapping of *MUTYH* variant effects using DNA repair reporters

**DOI:** 10.1101/2025.03.01.640912

**Authors:** Shelby L. Hemker, Ashley Marsh, Felicia Hernandez, Elena Glick, Grace Clark, Alyssa Bashir, Krystal Jiang, Jacob O. Kitzman

## Abstract

Variants of uncertain significance (VUS) limit the actionability of genetic testing. A prominent example is *MUTYH*, a base excision repair factor associated with polyposis and colorectal cancer, which has a pathogenic variant carrier rate approaching 1 in 50 individuals in some populations. To systematically interrogate variant function in *MUTYH*, we coupled deep mutational scanning with a DNA repair reporter containing its lesion substrate, 8OG:A. Our variant-to-function map covers >97% of all possible *MUTYH* point variants (n=10,941) and achieves 100% accuracy classifying the pathogenicity of known clinical variants (n=247). Leveraging a large clinical registry, we observe significant associations with colorectal polyps and cancer, with more severely impaired missense variants conferring greater risk. We recapitulate known functional differences between pathogenic founder alleles, and highlight sites of complete missense intolerance, including residues that intercalate DNA and coordinate essential Zn^2+^ or Fe-S clusters. This map provides a resource to resolve the 1,032 existing missense VUS and 90 variants with conflicting interpretations in *MUTYH*, and demonstrates a scalable strategy to interrogate other clinically relevant DNA repair factors.

## Introduction

The broad availability of clinical genetic testing has intensified the challenge of interpreting human genetic variation. It is particularly difficult to ascribe pathogenicity to missense variants because they vary widely in severity and mechanism, and most are individually rare. Consequently, many remain as variants of uncertain significance (VUS), contributing to anxiety for patients and families and lost opportunities for prevention and early detection.

Inherited cancer risk disorders undergo particularly widespread genetic screening and carry an accordingly heavy burden of VUS. One prominent example is *MUTYH*-associated polyposis (MAP), first described in families presenting with the characteristic polyposis phenotype of multiple colorectal polyps and carcinomas, but lacking inherited pathogenic *APC* variants^1,2^. Without intervention, colorectal cancer prevalence reaches 80% by age 70 in MAP, compared to 2-3% among the general population^3^. MAP also confers elevated risk for some extracolonic cancers^4–6^.

MAP is an autosomal recessive condition caused by inherited loss of function variants in *MUTYH*, the human ortholog of the DNA glycosylase *mutY*. This deeply conserved base excision repair factor limits the mutagenic potential of 7,8-dihydro-8-oxoguanine (8OG), the major byproduct of oxidative damage^7^. This lesion is mutagenic because of its tendency to mispair with adenine, which leads to a G:C>T:A transversion after replication (**Fig. 1A**). Tumorigenesis in MAP is often driven by somatic *KRAS* or *APC* mutations resulting from these transversions^8^, and individuals with MAP bear elevated genome-wide mutational signatures of oxidative damage both in normal tissue and tumors^9^.

**Figure 1.**
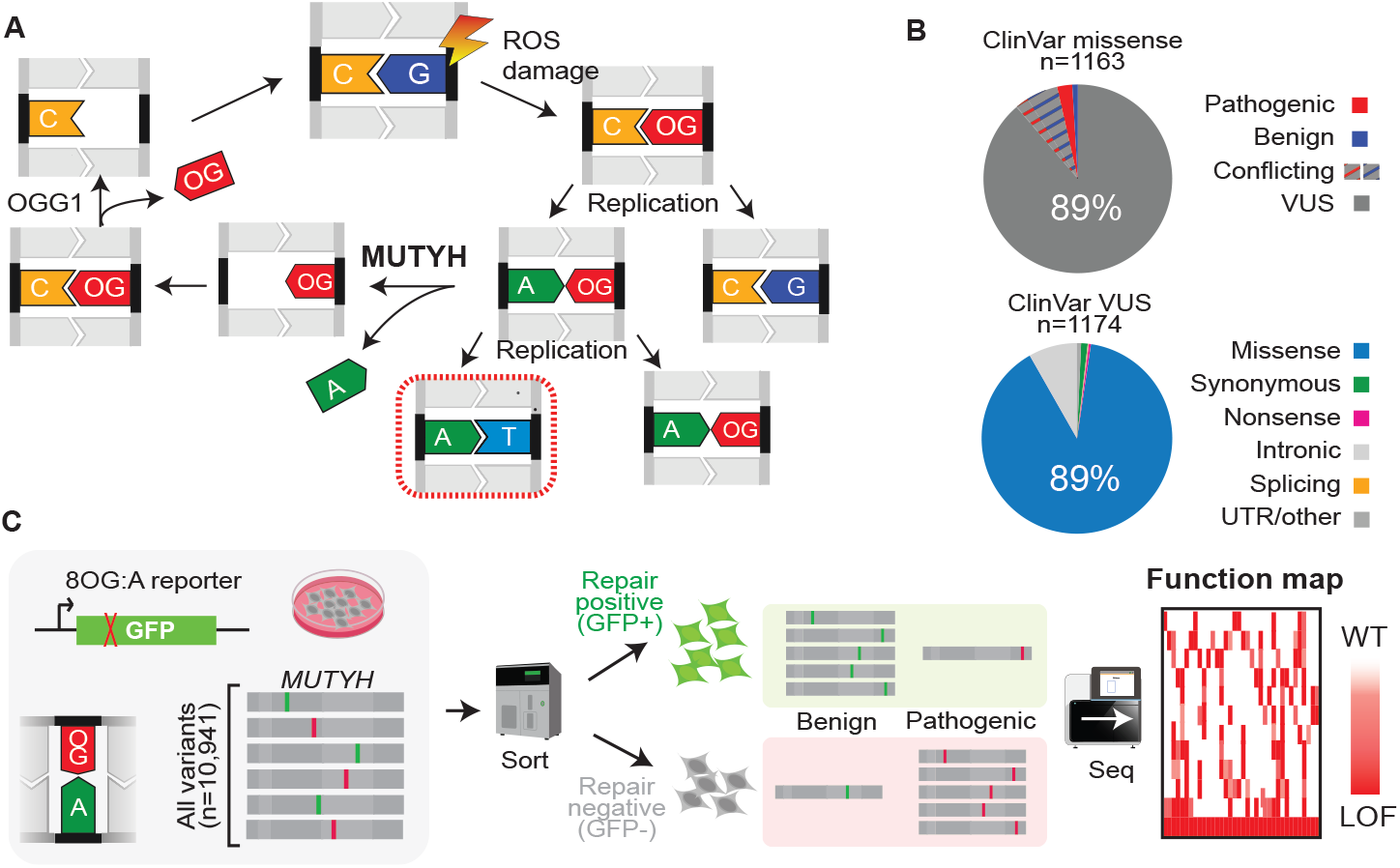
Overview of pooled variant-to-function mapping in *MUTYH*. **(A)** 7,8-dihydro-8-oxoguanine (OG) is a byproduct of oxidative damage and can mispair with adenine, leading to C:G → A:T transversion after successive rounds of replication without repair. MUTYH excises adenine opposite OG, followed by subsequent base excision repair steps. **(B)** Burden of clinical missense variants of uncertain significance (VUS) in *MUTYH*. Nearly all missense clinical variants in *MUTYH* are VUS or have conflicting reports (upper), and most *MUTYH* VUS are missense (lower). **(C)** Schematic of pooled functional assay for *MUTYH. MUTYH* mutant library cell population is transfected with an OG:A-bearing reporter, which expresses GFP only after proper repair. Cells with functionally intact, benign *MUTYH* variants are sorted to the GFP+ bin, while those with non-functional pathogenic variants are sorted into the GFP-bin. *MUTYH* mutations are quantified within each sorted population by sequencing and scored by their relative enrichment in GFP+ versus GFP-to yield a variant-to-function map.

Genetic screening for MAP can be lifesaving because, with proper surveillance, progression to cancer can be prevented^10^. However, personal and family history alone often fail to identify affected individuals due to the recessive inheritance pattern, incomplete penetrance, and variability in polyp number and age of onset^11–13^. Several founder alleles are well-recognized, including c.452A>G (Y151C) and c.1103G>A (G368D), for which nearly 2% of individuals in European ancestry groups are heterozygous carriers^14,15^. However, a long tail of individually rare variants poses an interpretation challenge: of the 1,163 *MUTYH* missense variants listed in the ClinVar database^16^, 89% are VUS (**Fig. 1B**), and over half of these are so rare as to be unobserved in exome sequencing databases covering over a million individuals^15,17^.

Functional evidence can guide variant interpretation, and various assays have been devised to test *MUTYH* variants. The normal function of MUTYH is to remove adenine opposite 8OG, with subsequent steps coordinated by other factors to remove and replace the oxidized base. *In vitro* reconstitution studies^1,18,19^ have provided valuable insights into the kinetics and remarkable specificity of this reaction, in which 8OG is distinguished from undamaged guanines present at ∼10^6^-fold molar excess^20^. However, purifying individual variant proteins is extremely laborious and suffers from batch variability. Fluctuation assays can test variants’ effects on mutation rate, but have been implemented only in bacterial hosts^21,22^, while small numbers of variants have been tested in mammalian cells using reporters that couple 8OG repair to fluorescent protein expression^23,24^.

Despite contributing key mechanistic insights, single-variant studies inevitably provide an incomplete picture, and do not readily scale to meet the current VUS burden. Here, we couple an optimized DNA repair reporter with saturation mutagenesis to systematically measure variant effects across *MUTYH*, yielding a variant-to-function map of nearly every possible single-residue variant (97.7% of n=10,941; **Fig. 1C**). These scores perfectly separate pathogenic and benign clinical variants with existing high-confidence classifications. Overlaid on a clinical laboratory database with individual-level records, they are strongly predictive of MAP phenotypes including polyps and early-onset colorectal cancer. A large fraction of missense variants exhibit intermediate functional defects, including some with strong clinical evidence for pathogenicity; these alleles may be sensitive to modifier effects^25,26^. This resource will guide the resolution of the >1,100 standing *MUTYH* missense VUS and enable prospective interpretation of newly detected rare variants, improving the accuracy and equitability of genetic testing.

## Results

### A scalable reporter of 8OG:A repair

To enable high-throughput measurement of *MUTYH* variant function, we leveraged a repair reporter assay^23,24^, in which a single 8OG:A mispair results in early truncation of a GFP expression cassette. Proper repair restores the open reading frame and with it, fluorescence **(Fig. 2A)**. With existing approaches it may be challenging to obtain the quantity and purity of reporter needed for large-scale transfections, so we devised a novel strategy leveraging enzymatic ssDNA production^27^ with a single 8OG incorporated by *Taq* DNA polymerase (**Supplementary Fig. 1**). We verified efficient, site-specific incorporation of 8OG by Sanger sequencing, with stalling observed as the polymerase encountered the lesion, followed by mixed (∼2:1) incorporation of A and C (**Fig. 2B)**.

**Figure 2:**
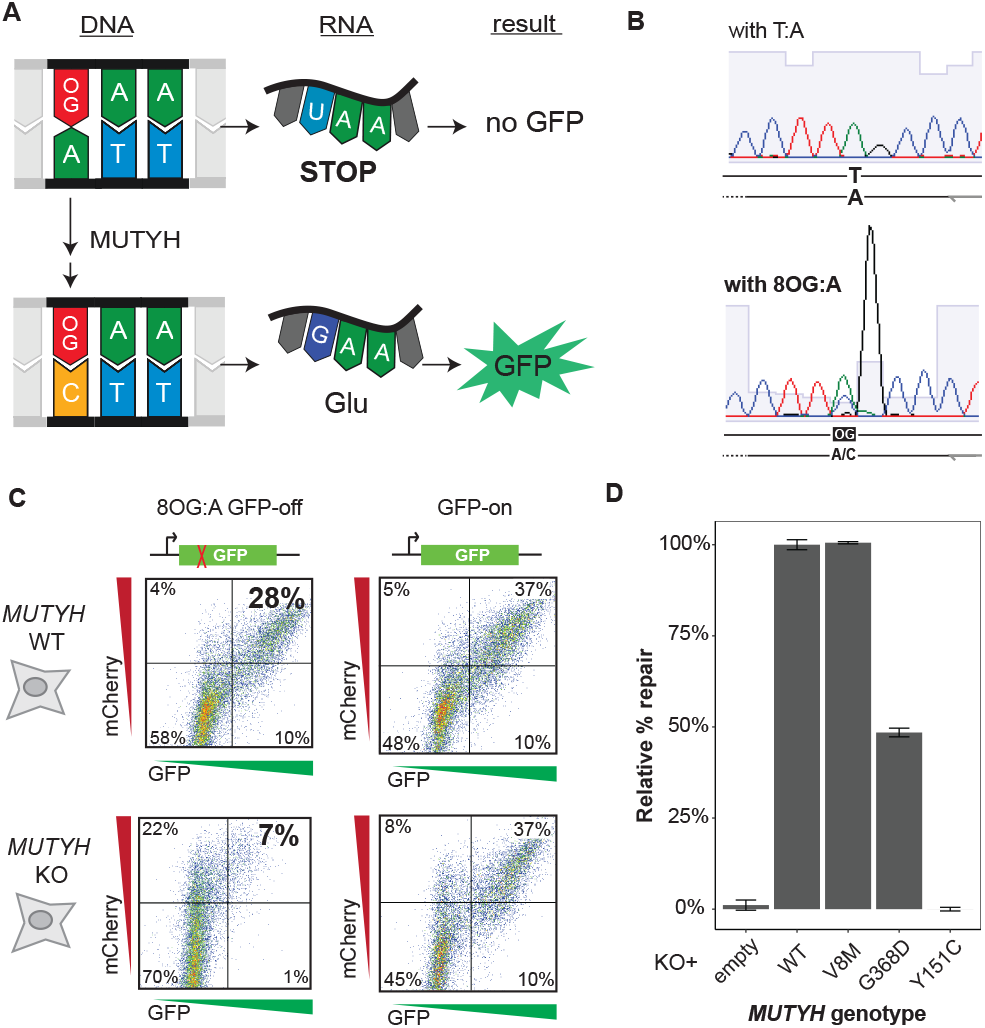
Functional reporter for MUTYH activity. **(A)** GFP expression construct engineered to contain an 8OG:A mismatch at codon 35. The transcribed strand contains a stop codon, but excision of adenine by MUTYH and subsequent replacement by cytosine restores a glutamine codon and GFP expression. **(B)** Validation of site-specific 8OG incorporation by Sanger sequencing. Compared to unmodified T:A (upper), the 8OG:A construct (lower) causes stalling as the polymerase encounters 8OG, followed by mixed A/C incorporation. **(C)** Functional validation of 8OG:A reporter, by FACS for GFP (repair signal) and mCherry (included as a transfection efficiency control). Parental HEK-293 cells (upper row) have four-fold higher repair-positive rate compared to *MUTYH* KO cells (lower row). A construct without 8OG or stop codon (‘GFP-on’; right column) indicates similar transfection efficiency between the two lines. **(D)**. Low repair activity in KO cells is restored after complementation with WT *MUTYH* or V8M, a common, benign polymorphism. Cells stably expressing pathogenic founder mutations G368D and Y151C show partial or complete loss of MUTYH activity, respectively. Repair percentage is quantified as (%GFP+)/(%mCherry+), and normalized to WT-transduced cells. Error bars, s.e.

We transfected this reporter into HEK-293 cells and an isogenic clonal *MUTYH* knockout (KO) line. We used flow cytometry to quantify the percent of cells positive for both GFP, indicating A→C repair opposite 8OG, and mCherry, included to control for transfection efficiency. Repair levels in *MUTYH* KO cells were reduced by 76% compared to wild-type parental cells (**Fig. 2C, Supplementary Fig. 2**), while GFP expression was unchanged using a reporter bearing an intact open reading frame (unmodified G:C instead of 8OG:A), indicating that 8OG:A repair is compromised in the absence of MUTYH.

We next verified that this reporter could distinguish known benign and pathogenic *MUTYH* missense variants (**Fig. 2D**). Whereas reintroducing WT *MUTYH* or the known benign polymorphism V8M each robustly restored repair, the pathogenic founder variant Y151C showed no activity above KO cells’ background levels. Another, G368D, was reproducibly intermediate (48% activity), consistent with purified protein studies^1^, and with both the lower mutational burden^9,28^ and later clinical age of onset^12,29^ in individuals homozygous for G368D versus Y151C. Thus, this reporter quantitatively measures *MUTYH* variant function across a range of activity.

### A variant-to-function map of *MUTYH*

To expand this assay to systematically interrogate *MUTYH* variants, we performed saturation mutagenesis^30^ of the predominant, 521-codon nuclear isoform cDNA^31^. We obtained near-complete coverage of the single-codon mutational space, with 99.0% of intended variants present at a frequency of ≥1/10,000 (**Supplementary Fig. 3**). These libraries were stably integrated in HEK-293 *MUTYH* KO cells at low titer so that each resulting cell expressed a single *MUTYH* variant. Mixed mutant cell pools were co-transfected with the 8OG-GFP reporter and mCherry constructs, then sorted to separate cells displaying repair activity (mCherry+, GFP+) from those positive for transfection but with low repair signal (mCherry+, GFP-). From the pre-sorting cell pools as well as each sorted population, the stably integrated mutant *MUTYH* cDNA library was subjected to deep amplicon sequencing.

As expected, repair-positive sorted cells were enriched for synonymous variants and depleted for nonsense variants, while repair-negative sorts showed the reciprocal pattern (**Supplementary Fig. 4**), demonstrating the feasibility of pooled selection on *MUTYH* functional status. Using Rosace^32^, we calculated a log_2_-scaled function score quantifying variants’ enrichment or depletion in the repair-positive vs the repair-negative sorts.

The resulting variant-to-function map provides measurements for 98% of the possible single-codon variants in *MUTYH*, and broadly reflects missense constraint at known structural domains and key residues (**Fig. 3; Supplementary Tables 1 and 2**). Function scores perfectly separated synonymous and nonsense variants across codons 1-471 (n=725; area under the precision-recall curve, prAUC=1; **Supplementary Fig. 5**). Nonsense variants at 472 onward were uniformly tolerated (mean score: -0.03), indicating this disordered C-terminal region is dispensable for 8OG:A repair. Though this assay does not model the effects of nonsense-mediated decay (NMD), these codons all reside in the final exon of *MUTYH* or within 55 nt of the 3’-most splice site and as such are likely to escape NMD *in vivo*.

**Figure 3:**
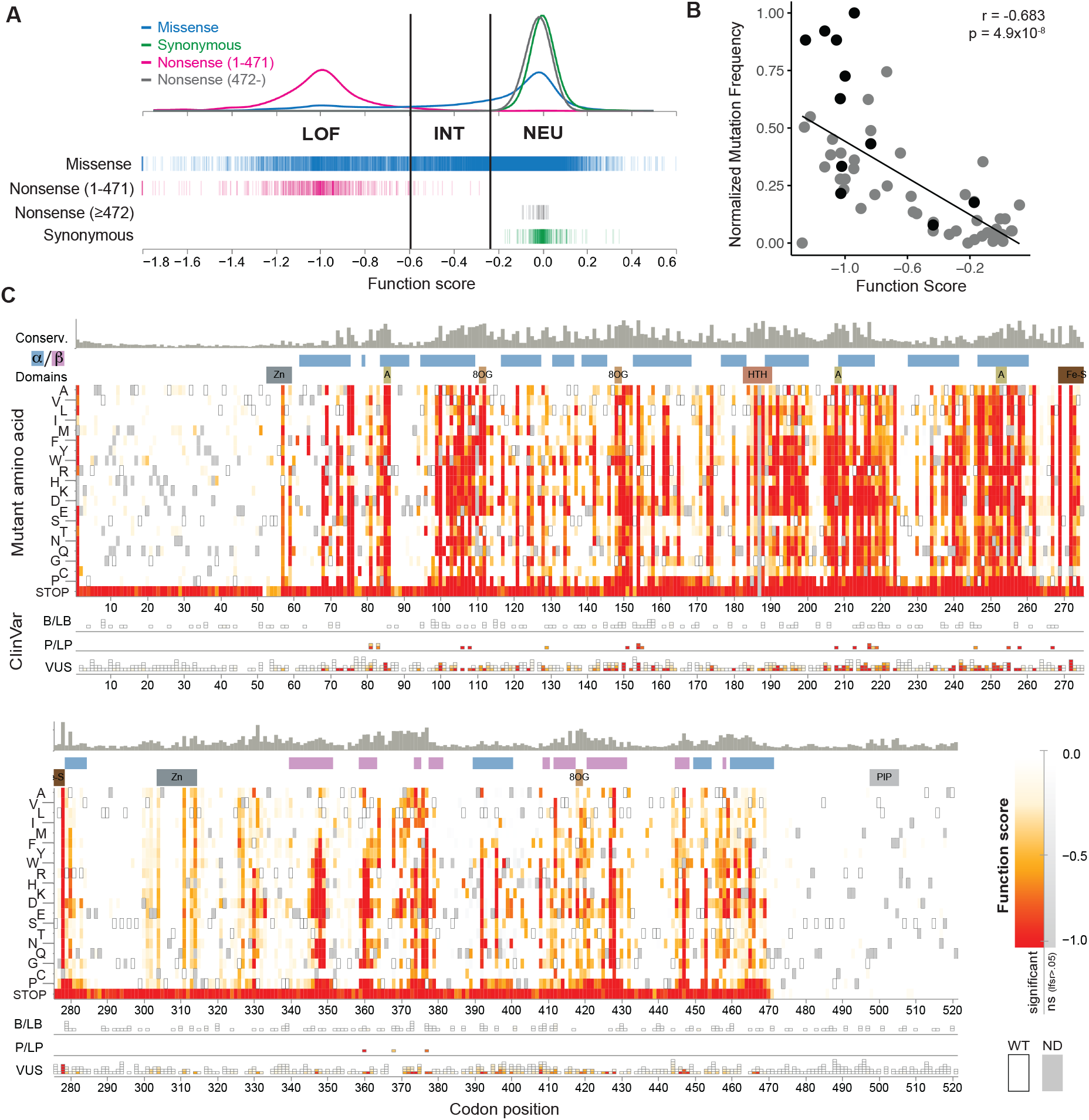
A complete variant-to-function map across *MUTYH*. **(A)** Distributions of function scores by variant class; nonsense variants in codons 1-471 are perfectly separated from synonymous variants. Vertical lines denote cutoffs for function classifications (LOF: loss of function, INT: intermediate, NEU, neutral). **(B)** Validation of pooled assay function scores is shown by strong correlation to relative mutation rates as previously measured by fluctuation tests (n=50 *MUTYH* missense variants; gray points: ref ^22^, black: ref. ^21^). Mutation rates are rescaled relative to wild-type and knockout *MUTYH* as measured within each prior study. **(C)** Variant-to-function heatmap across *MUTYH*, shaded by function score (white: WT-like; red: null-like) for each mutant amino acid (rows) at each codon position (columns). Mutations that do not pass multiple testing correction (lfsr>0.05) are shaded from white to gray, wild-type residues are boxed, and dark gray denotes missing data. Tracks above heatmaps, from top to bottom: evolutionary conservation score, protein secondary structure, key domains. Below heatmaps, ClinVar variants are shown in stacks, grouped by interpretation (B/LB: benign/likely benign; P/LP: pathogenic/likely pathogenic; VUS: uncertain or conflicting), and shaded as in the heatmap.

Missense variants spanned the full score range, underscoring that their functional defects are continuous in nature, rather than all-or-none. Scores were broadly consistent with previous functional studies, showing strong correlation with mutation rates from *E. coli*-based fluctuation assays of 50 human *MUTYH* missense variants^21,22^ (**Fig. 3B**). Using cutoffs based upon nonsense and synonymous score distributions (**Methods**), we classified 9,776 missense variants as loss-of-function (LOF), intermediate (INT), or neutral (NEU). Excluding the unstructured termini (codons 1-51 and 472-521), 25% of missense variants scored as LOF and 59% scored as NEU, leaving a substantial proportion (16%) as intermediate (INT). Among these was the founder allele G368D, consistent with individual variant assays (**Fig. 2D**), as well as several others (P129L, V206M, V218F) previously shown to retain partial activity^21^. These examples demonstrate that this pooled 8OG:A reporter assay retains the dynamic range to distinguish between different degrees of functional impairment.

### Accurate classification of clinical missense variants

To evaluate these scores’ potential as functional evidence, we asked whether they could recapitulate existing clinical variant interpretations. To obtain a ‘truth set’ of variants with confident interpretations, we stringently filtered *MUTYH* SNVs from ClinVar, requiring at least two clinical submissions per variant, and excluding any with VUS/conflicting interpretations or likely splicing defects (SpliceAI^33^ deltaMax score≥0.3). This resulted in a set of 247 variants (**Fig. 4A)**, with 187 classified as pathogenic/likely pathogenic (P/LP) and 60 as benign/likely benign (B/LB). Our function scores perfectly separated these sets (prAUC=1, **Fig. 4B**), and applying score cutoffs (treating LOF and INT as pathogenic and NEU as benign) resulted in 100% concordance with the standing classifications. Performance was nearly as high on a less stringently filtered set allowing for conflicting submissions from different clinical laboratories (n=460, prAUC=0.998; **Supplementary Fig. 5**). Thus, our variant-to-function map supports *MUTYH* variant classification with near-perfect specificity and sensitivity.

**Figure 4:**
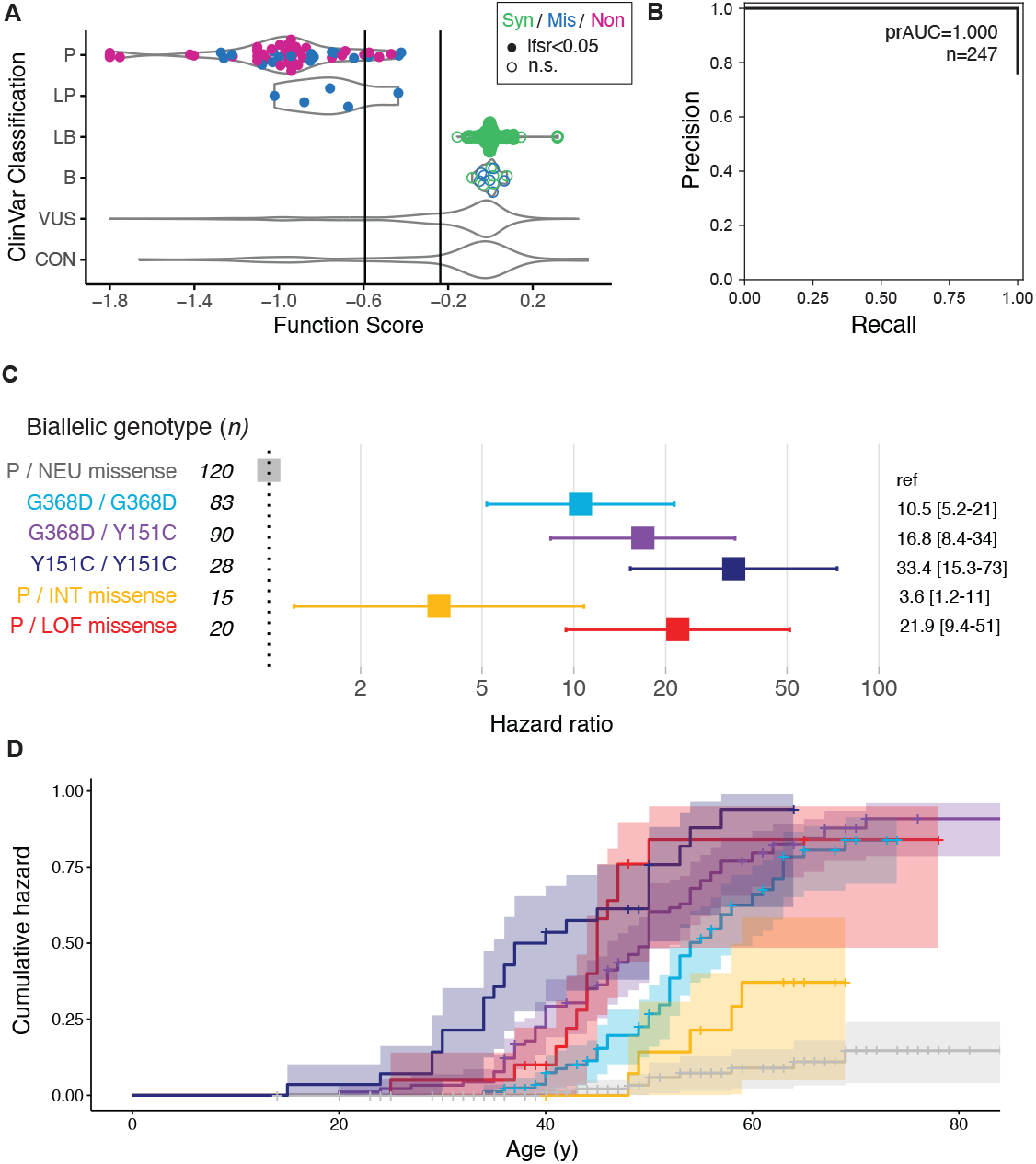
Accurate classification of *MUTYH* clinical variants conferring MAP risk. **(A)** Function scores perfectly separate pathogenic from benign variants among a high-confidence set of 247 SNVs from ClinVar stringently filtered to exclude low-confidence or singleton submissions. Point color denotes variant type, and filled/open points denote statistical significance. Variants with uncertain (VUS) or conflicting (CON) interpretations in ClinVar are shown as violin plots. **(B)** Precision-recall curve showing classification performance between P/LP and B/LB variants in (A). **(C)** Cox proportional hazards modeling of MAP phenotypes (polyps or early onset colorectal cancer) among 356 individuals biallelic for *MUTYH* variants. Individuals carrying one known pathogenic variant and one neutral-scoring missense VUS are the baseline group (gray), while genotypes comprising two pathogenic founder variants, or one pathogenic variant plus one missense INT or LOF VUS as measured by pooled assay, all confer significant hazard ratios. **(D)** Survival curves showing cumulative hazard for MAP phenotype by age, separated by biallelic genotype (curves shaded by genotype as in (C)).

We further scrutinized assay performance on missense variants, given the interpretation challenge they pose. We leveraged a clinical laboratory database of *MUTYH* variant carriers, for which individual-level genotype, demographic, and clinical records were available, unlike in public-facing databases such as ClinVar. This allowed us to exclude variants whose classifications relied upon prior functional assays, avoiding the circular logic of validating one functional assay with a previous one. This left 87 missense variants previously classified as benign or pathogenic, of which our map agreed with 86 (100% sensitivity, 98% specificity), underscoring that classification performance was not limited to distinguishing silent from truncating mutations.

To quantify the strength of evidence provided by our map, we used the framework implemented by the ClinGen Sequence Variant Interpretation group^34^, which incorporates the number of variants in the truth set as well as assay classification performance. Using the stringently filtered truth set (n=247), an abnormal function score call results in an OddsPath (odds of pathogenicity) of 187. Conversely, the OddsPath for a neutral assay result was 0.0167 (1/59.9). Both of these correspond to ‘strong’ evidence codes (PS3/BS3) under the ACMG variant interpretation framework, as did scores derived from the clinical missense-only set (**Table 1**).

**Table 1.** Functional assay calibration using *MUTYH* clinical variants.

| Truth set | n, total | Pathogenic variants |  |  |  |  | Benign variants |  |  |  |  |
| --- | --- | --- | --- | --- | --- | --- | --- | --- | --- | --- | --- |
|  |  | n | Abnormal (LOF/INT) | Normal (NEU) | Odds Path | Evidence code | n | Abnormal (LOF/INT) | Normal (NEU) | Odds Path | Evidence code |
| **ClinVar (stringent)** | **247** | 60 | 60 | 0 | 187 | PS3 | 187 | 0 | 187 | 1/59.9 | BS3 |
| **ClinVar (lenient)** | **460** | 94 | 91 | 3* | 354 | PS3_vs | 366 | 0 | 366 | 1/31.3 | BS3 |
| **Missense (Ambry)** | **36** | 36 | 36 | 0 | 51 | PS3 | 51 | 1 | 50 | 1/35.3 | BS3 |

Sets of known pathogenic or benign variants were used to quantify the strength of evidence provided by assay function scores. Three truth sets consisted of all *MUTYH* SNVs listed in ClinVar with conflicting or unsupported entries alternatively removed or retained (stringent and lenient filtering; see Methods), and a set of exclusively missense variants from a single clinical laboratory supported by individual-level clinical information. OddsPath scores were calculated from the counts of variants with concordant/discordant functional results<sup>34</sup>, with corresponding strength-evidence codes (PS3: strong evidence for pathogenicity, PS3\_vs: very strong evidence for pathogenicity, BS3: strong evidence for benignity). \*Note: 2 of 3 discordant pathogenic variants in the ClinVar lenient truth set are C-terminal truncating variants (codons 472 and 473).

### Missense loss of function confers MAP risk

Using individual-level clinical records, we estimated the risk conferred by *MUTYH* VUS. Our clinical database contains 155 individuals who each carry both a missense VUS and a known pathogenic variant (primarily one of the two founder missense alleles; **Supplementary Table 3**). These individuals’ risk would be greatly elevated if their VUS were pathogenic. To test this, we asked if the burden of MAP-associated phenotypes differed based upon the VUS’ functional assay results. Of these individuals, 104 were phenotypically informative, having either CRC by age ≤50, or polyp count ≥10 at any age, or conversely, having reached age ≥60 without personal history of polyps or CRC. We observed a significant excess of MAP phenotypes in individuals who carried LOF versus NEU missense VUS alongside a known pathogenic variant (**Fig. 4C**; hazard ratio=21.9 [95%CI: 9.4-51.0], *p*=7.7×10^−13^, Cox proportional hazards test). Intermediate-scoring missense VUS were associated with milder but still significant risk (HR=3.6 [1.2-10.8], *p*=0.02).

For context, we compared these results to the 201 individuals who carried known risk genotypes: homozygous for either founder missense variant, or compound heterozygous for each. Observed MAP risks recapitulated known genotypic differences: Y151C homozygotes had the highest risk, followed by compound heterozygotes and then G368D homozygotes (**Fig. 4D**), comparable to population-based estimates from UKB^35^. Compared to these, functionally abnormal missense VUS conferred risk most similar to Y151C/G368D compound heterozygosity.

### Systematic, ancestry-agnostic VUS resolution

We next set out to reclassify standing missense VUS. Our map covers 99.4% of the 1,122 missense *MUTYH* SNVs with VUS or conflicting interpretations listed in ClinVar. Of these, 26% had an abnormal functional assay result (LOF: 14%, INT: 12%), with the rest either normal (71%), or neutral at the protein level but predicted as splice-disruptive (2%). These 295 clinical LOF or INT VUS have the potential to be reclassified to likely pathogenic with the addition of this functional evidence. Notably, in our clinical database, six of these functionally abnormal VUS were observed co-occurring with known pathogenic variants in individuals who did not yet display a MAP phenotype. Given that these individuals are younger (median: 40 years) than the typical age of onset, and that in some cases, CRC may develop in the absence of polyposis^6^, reclassifying their VUS provides the opportunity for prevention or early intervention.

Pathogenic *MUTYH* variants are routinely found in population databases, consistent with MAP’s recessive inheritance (**Fig. 5**). Excluding Y151C and G368D, the 169 other functionally abnormal missense variants found in gnomAD are individually rare (median MAF: 1.4×10^−6^). However, with a cumulative MAF of 0.17%, their overall contribution is comparable to founder alleles’. Some of these may remain unclassified due to biases in clinical sequencing and variant interpretation^36,37^. Using ancestry-specific allele frequencies from gnomAD^15^ and RGC-ME^17^, we identified 318 *MUTYH* missense SNVs with at least 5-fold higher frequency in a non-European versus the primary European ancestry groups in each database. Among these, 84 were functionally abnormal, with the great majority either absent from ClinVar (23%) or having VUS/conflicting interpretations (64%), despite being on average 220-fold more common in their respective ancestry groups.

**Figure 5:**
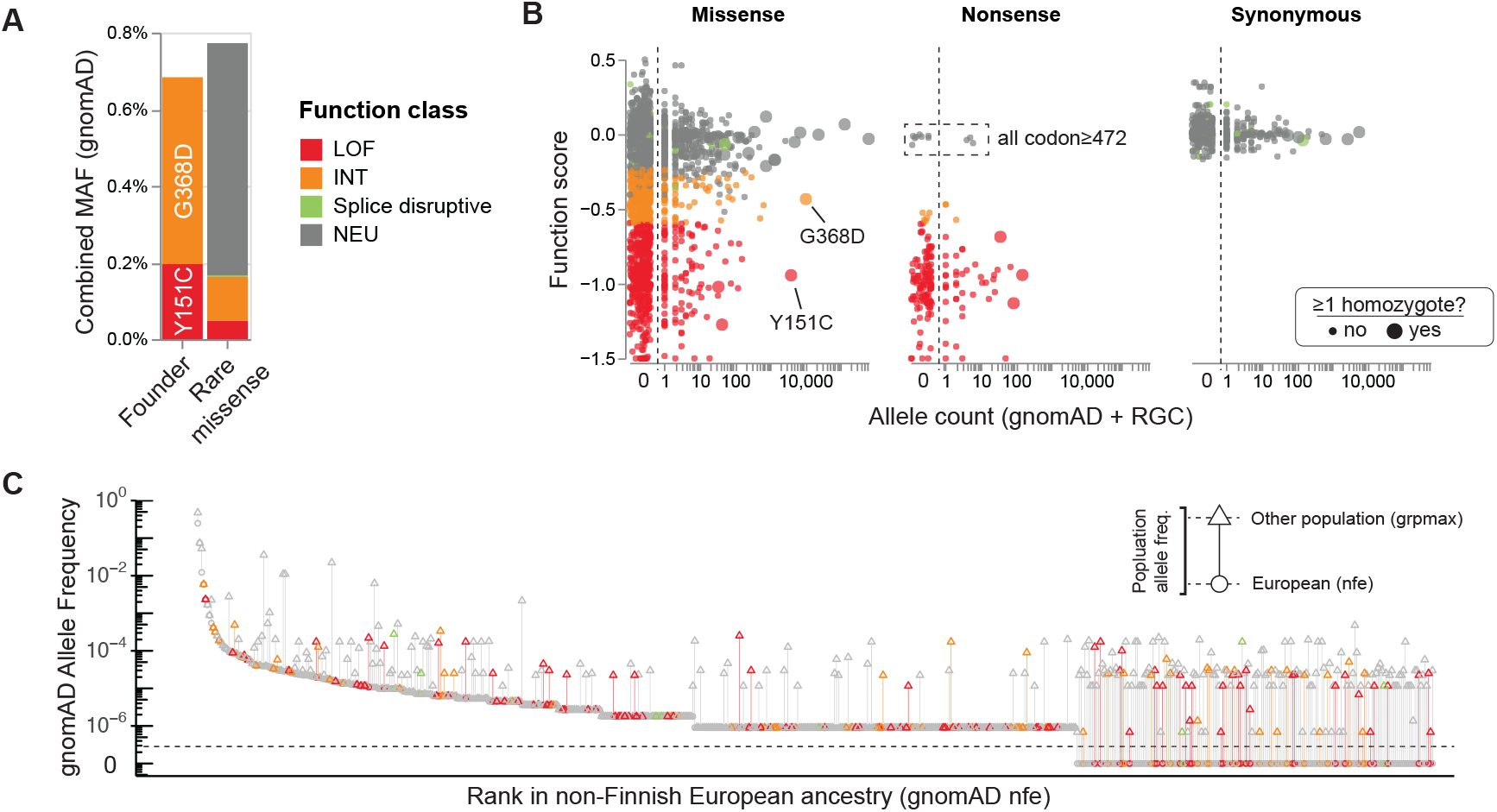
SNVs in population databases. **(A)** Comparable cumulative load of founder alleles (G368D, Y151C) and rare missense variants in gnomAD (popmax MAF≤0.75%). Rare variants are shaded by functional assay classification. **(B)** Function scores versus population allele frequency (sum of gnomAD non-UKB and RGC allele counts), by variant class. Larger points correspond to variants observed in the homozygous state. Neutral scoring nonsense variants are exclusive to C-terminus (codon≥472). **(C)** Functional status of rare missense variants in different populations. *MUTYH* SNVs from gnomAD are shown, with minor allele frequency plotted versus allele frequency rank in the most deeply ascertained population (nfe, non-Finnish European). Variants that are more common in other populations are shown with a vertical line extending from the nfe minor allele frequency (circle) to the maximal per-population allele frequency (triangle). Variants observed exclusively outside of nfe are below the dotted line.

### Structure-function relationships in MUTYH

We next compared our variant-function map with structural features of MUTYH. We used an AlphaFold3-predicted^38^ structure of human MUTYH in complex with 8OG:A which aligned closely with experimentally resolved crystal structures of the mouse and human proteins^22,39^ (**Supplementary Fig. 6**). As expected, LOF and INT variants were heavily enriched among buried versus exposed residues (OR=35.7 and OR=6.6, respectively; Fisher’s exact *P*=10^−185^ for both; **Fig. 6A**). Changes in folding free energy predicted by FoldX^40^ correlated modestly with function score (Pearson’s r=-0.52), with LOF being more highly destabilizing than INT variants (median difference 1.99 kcal/mol; *p=*6.9×10^−65^, Mann-Whitney U), which were in turn more so than NEU variants (1.80 kcal/mol, *p*=1.3×10^−185^). These results are consistent with protein destabilization accounting for the dysfunction of many, but not all, missense variants^41^.

**Figure 6:**
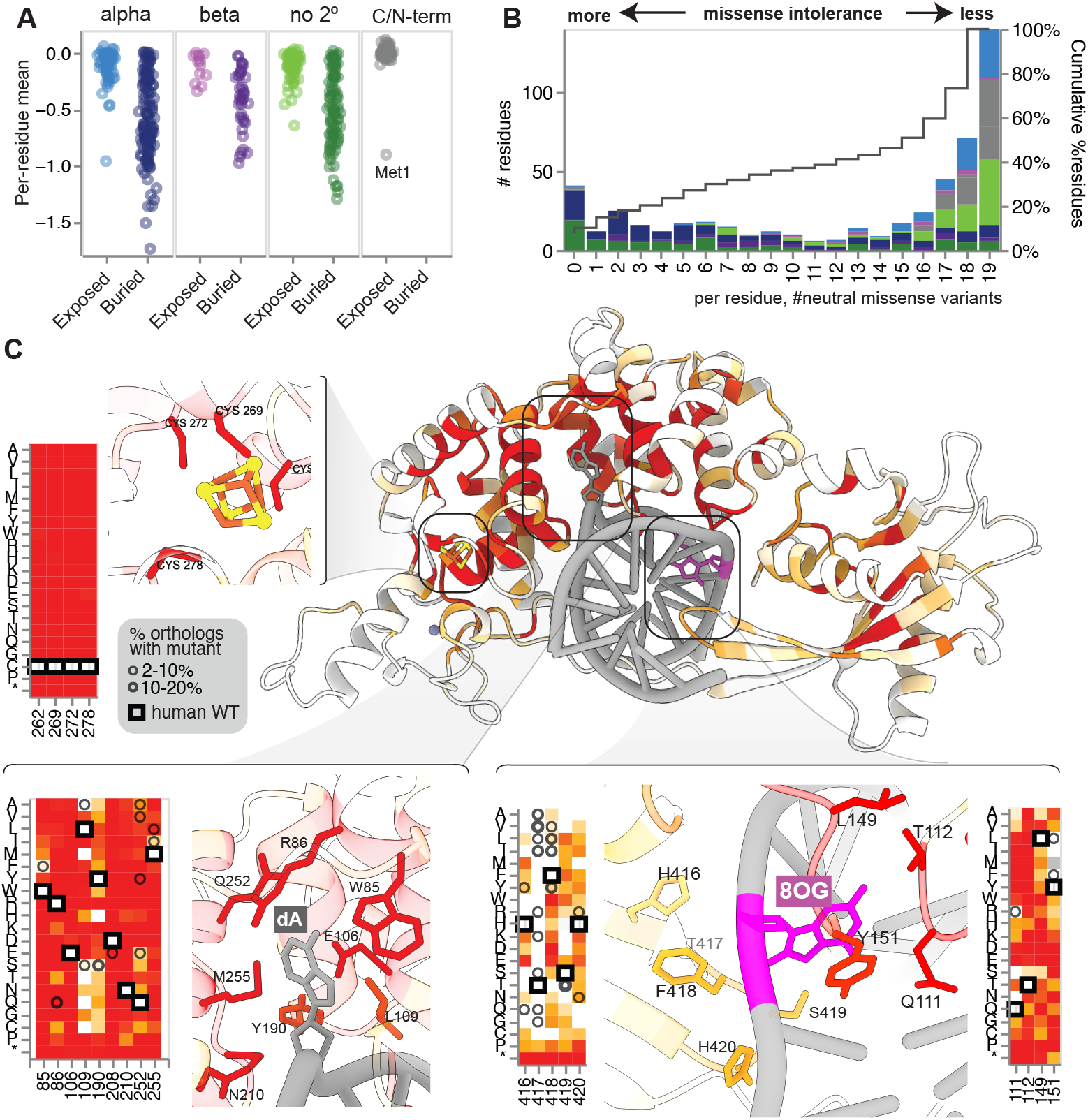
Missense constraint shapes structure-function relationships. **(A)** Per-residue mean missense function scores, grouped by residue surface area exposure status and secondary structure. Buried residues are enriched for deleterious variants. **(B)** Distribution of per-residue constraint across MUTYH, with a histogram showing the number of mutations per residue which are functionally neutral (ranging from 0 to 19, indicating complete intolerance and tolerance to missense mutations, respectively). Counts are shaded by residues’ structural properties, as in (A). **(C)** AlphaFold3 predicted structure for MUTYH bound to 8OG:A pair (purple/gray). MUTYH residues are shaded by mean missense function score (shaded as in Figure 3C; darker red indicates more severe). Key functional regions are shown in insets with corresponding variant-function map details. Counter-clockwise from upper left, [4Fe-4S] cluster, adenine-binding/catalytic site, HxFSH loop/8OG-contacting residues. Within heatmaps, wild-type residues are boxed, and circles denote residues observed in MUTYH orthologs.

A substantial minority of residues were either fully missense-intolerant (7.9%), or nearly so (15.0% with ≤2 of 19 possible missense tolerated as NEU; **Fig. 6B**). Overlaid on the MUTYH structure (**Fig. 6C**), this constraint highlighted key functional sites, including at the essential [4Fe-4S] cluster which mediates lesion engagement and recognition^22,42,43^. As expected, mutations at any of the four coordinating cysteines resulted in LOF comparable to truncation (mean score: -0.90). Similarly complete constraint was also observed at residues known to be critical for recognition and removal of the extra-helical ‘flipped out’ adenine^39,44^ including the catalytic residues D208 and E106 as well as an allosteric network (including N210, R213, and R217) connecting these sites to the [4Fe-4S] cluster^22^. Thus, strong constraint highlights the most functionally sensitive sites in MUTYH, in and around the catalytic center.

A similar but weaker pattern of constraint was seen at the ‘zinc linchpin’ motif^45^ which plays a role in orienting the catalytic and lesion recognition domains (**Supplementary Fig. 7**). At this site, a Zn^2+^ ion is coordinated by four residues, with both C216 and H57 nominated as the fourth^39,46^. Our data support a primary role for H57, as it is intolerant to all missense mutation, whereas C216 tolerates mutations to Thr, Ser, and Ala. Missense effects at these four Zn-coordinating residues (H57, C304, C311, C314) were milder than those at the [4Fe-4S] cluster (mean: -0.55 vs -0.90).

Residues involved in 8OG lesion recognition showed a more nuanced pattern of constraint (**Fig. 6C**). At the key ‘probe’ residue Y151, which intercalates into the double helix near 8OG, 10 of 19 mutations scored as LOF, including the founder allele Y151C. However, in addition to the wild-type Tyr, other bulky hydrophobic residues (Phe, Leu, Ile, Met) were tolerated. Consistent with this pattern, some DNA glycosylases have Phe or Leu at the equivalent position^7,47^, and *E. coli mutY* mutants with these residues retain WT-like activity^48^. Mutations at Q111, T112, and L149, which initiate 8OG:A recognition via contacts with the 2’-amino group, had severe effects (42/57 scored as LOF). Opposite this, near the 8OG 7’NH, mutations at the conserved HxFSH motif exhibited intermediate effects, aside from the non-conserved second (‘x’) position which tolerated all mutations except proline. Relaxed constraint at the HxFSH motif is consistent with a secondary role in human MUTYH as compared its essentiality in bacterial *mutY* orthologs^49,50^.

Certain substitutions were systematically disruptive, such as proline mutations within secondary structures (mean score: -0.68). However, a subtler ‘ribbon’ of constraint against negatively charged aspartic and glutamic acid mutations was also evident. These included the founder allele p.G368D, which introduces a negative charge ∼3.5 Å from the DNA backbone at the 8OG lesion-adjacent base^7,42,51^. A similar pattern held throughout the 50 residues within 5A of the backbone, where mutations to D or E were significantly more disruptive than others (median score difference from D/E versus other amino acids among backbone-proximal residues: -0.20, vs -0.09 among backbone-distal residues; *p*=4.8×10^−4^ one-sided MW U test; **Supplementary Fig. 8**). The signal of constraint was specific to D/E: of the 153 pairs of amino acids, it was the only disfavored pair (Bonferroni-corrected p>0.10 for all other pairs), and it persisted after controlling for backbone-proximal residues’ surface exposure (*p*=7.7×10^−3^, permutation test). Thus, DNA backbone interactions broadly contribute to the landscape of missense constraint across MUTYH.

Lastly, we asked whether variant effects could be predicted by structural and evolutionary features. Fitting logistic regression models of measured constraint to these features showed strong correlation overall (R^2^=0.56) but revealed 20 residues where the measured effects were much more severe than predicted by computational models (**Supplementary Fig. 9**). Among these were V301 and E302, which are known to mediate interactions with the 9-1-1 DNA damage signaling complex^52,53^. Another completely missense-intolerant residue, D224, resides in a small loop near the interdomain connector at the proposed APE1 interaction site^39^. Two others, Q240 and R246, similarly face outwards in or near unstructured regions. Mutations at these sites may disrupt important interactions such as the ‘baton pass’ of the abasic product from MUTYH to APE1 for downstream base excision repair^54,55^. These putative interaction sites may be poorly captured by intra-sequence structural or evolutionary coupling models.

## Discussion

Saturation mutagenic screens are rapidly contributing functional evidence^56–60^ which can inform clinical variant interpretation^61,62^. However, the underlying assays commonly rely upon cellular proliferation or drug sensitivity to separate mutations by function. Consequently, the ∼90% of genes which are dispensable for growth in culture^63^ remain inaccessible to these approaches and require custom approaches^64^.

In the current study, we systematically mutagenized the 10,941 single-residue variants in *MUTYH*, a prominent recessive polyposis disorder gene. Our approach combines, for the first time, deep mutational scanning with a direct readout of DNA damage repair from a lesion-bearing reporter. The accuracy of the resulting variant-to-function map is demonstrated by perfect separation of truncating and silent mutations. Between these extremes, missense mutations’ effects varied continuously, with the 25% impaired comparably to nonsense mutations classified as LOF, and another 16% displaying intermediate (INT) activity.

To validate these functional measurements for clinical use, we obtained high-confidence classifications from ClinVar and a large clinical laboratory database. We observed near-perfect concordance with our map, confirming its potential to serve as strong evidence for or against pathogenicity within the OddsPath framework^34^. We noted a minority of ClinVar entries, which did not meet criteria for inclusion in our truth sets, for which the listed interpretation was discordant with our functional classification (**Supplementary Table 4**). Most of these lacked submission criteria, were somatic mutations, or were incidental findings in non-MAP-associated tumor types, and their classifications should be reconsidered in light of this functional evidence.

Leveraging a clinical genetic testing database with detailed phenotypes, we found that variants with abnormal function in our assay strongly associated with polyposis and cancer. With 356 individuals biallelic for a known pathogenic *MUTYH* variant and a second rare variant – an order of magnitude more than found in UK Biobank^35^ – this result underscores the utility of intersecting MAVE data with high-volume clinical genetic testing databases. Therefore, the 1,122 *MUTYH* missense variants listed in ClinVar with a VUS or with conflicting interpretation can be re-evaluated with this functional evidence. Another 696 missense LOF/INT SNVs are as yet undetected in available clinical and population databases, indicating that sequencing has not yet saturated the repertoire of possible pathogenic missense variants in *MUTYH*. The resource provided by this study will enable their rapid classification as they are observed in the coming years.

Intermediate MAVE scores can be difficult to interpret, as they may reflect measurement noise or, instead, truly hypomorphic activity. Here, several lines of evidence suggest the latter. First, no synonymous variants are found in the INT range, indicating a low false positive rate. Secondly, we detect INT variants, including V218F and C304S, previously shown to have relatively mild effects in biochemical or mutation accumulation assays^21,39^. Finally, several clinical variants known to be partially defective score as INT, including V206M^21,65,66^ and the founder allele G368D, which is associated with delayed age of onset^12,67^ and lower mutation burden^9,28^.

The 16% of missense variants with intermediate function may be especially sensitized to modifier effects. These could arise from epistatic interactions from other 8OG pathway members: compound heterozygous loss of function for both *OGG1* and *MUTYH* results in ∼10-fold higher somatic mutation burden^9,68^, and similar effects are observed in mouse models^69,70^. Germline *de novo* C>A mutation rate is also increased^65,71^, an effect accentuated by subtle expression variation in other factors^26^ or *MUTYH* itself^25,72^. Intermediate missense variants’ penetrance for MAP phenotypes could be similarly shaped by *cis*-regulatory variation affecting *MUYTH* or its partners in BER.

Our map identifies functionally critical residues within MUTYH, particularly at the [4Fe-4S] cluster and at catalytic residues around the adenine, reflecting the dense interaction network nearby^22^. By contrast, mutations at the ‘zinc linchpin’ motif largely scored as INT (52%), with many having borderline-neutral scores. At these sites, our map defines an order of functional importance (H57>C311,C314>C304) consistent with prior mutation accumulation and *in vitro* catalysis measurements^39,45^, and with the presence of clinical likely pathogenic variants exclusively at H57. An excess of constraint nominates exposed residues D224, Q240, and R246 as potential protein-protein interaction sites; future studies will be required to confirm this and identify the interacting factor(s).

Systemically testing all single-codon residues in *MUTYH* highlighted patterns of constraint that would be inaccessible to SNV-only mutagenesis. For instance, W75 and Y76 have a strong preference for aromatics, while Y151 prefers bulky hydrophobic residues; but between these sites, only three of the ten tolerated amino acid substitutions can be reached with an SNV. Even at intensively studied sites such as Y151 and G368, our map demonstrates that assuming functional equivalence between different mutations at the same codon could lead to misclassification^73^.

There was a notable lack of constraint against C-terminal truncations from residue 472 onward. This unstructured domain is reported to interact with PCNA via a PIP box motif (residues 498-505), but this interaction appears to be dispensable for 8OG:A excision and subsequent repair in our reporter assay. Mutation accumulation studies in mESCs (a replication-coupled context) showed that a disruptive double-Phe-Ala mutation in the PIP box did not increase mutation rate, whereas the known pathogenic variant G368D did^74^. We cannot exclude a functional role for this region but note the clinical evidence is equivocal: in ClinVar, only seven truncating variants are listed among residues from 472 onward, nearly all from non-MAP individuals without a pathogenic second variant identified *in trans*. The lone apparent exception was a patient compound heterozygous for Q473* and G368D who presented with colorectal cancer and several adenomas. Given this contradictory evidence, follow-up studies are needed to define the C-terminus’ roles in genome stability and MAP.

Despite providing near-perfect classification accuracy on clinical variants, our study has several limitations, in particular that the cDNA-based approach cannot detect regulatory or splicing effects. Saturation editing strategies can capture these^75,76^, but usually at the expense of sparser mutational coverage, which can obscure mechanistically informative patterns of constraint. To identify splice-disruptive variants, we used SpliceAI^33^, a deep-learning splicing effect model which has been shown to be highly accurate by benchmarking against functional assays and clinical variants^77,78^; indeed, it almost perfectly identifies RNA-disruptive variants identified in saturation editing-based maps^59^. Nevertheless, endogenous locus editing has been shown as an effective approach in *MUTYH* and will provide a powerful toolset for deep characterization of individual variants of interest^79^.

In summary, we have shown how systematic variant-function mapping can assist with the classification of *MUTYH* clinical variants, while also providing insights into its structure-function relationships. As efforts to systematically map variant functional impacts across the human genome ramp up, our study provides a template to interrogate DNA damage response genes which underlie highly prevalent cancers and other disorders.

### Tables

## Methods

### Mammalian cell culture

Human HEK-293 cells (ATCC) were cultured in DMEM (Invitrogen) supplemented with 10% fetal bovine serum (Invitrogen) and 1% penicillin-streptomycin (Invitrogen). *MUTYH* knockout cells were derived by transfection with a Cas9/sgRNA plasmid (pSpCas9(BB)-2A-Puro, AddGene #62988) targeting *MUTYH* exon 4. Clonal knockout cells were isolated by limiting dilution and screened by PCR and sequencing (Plasmidsaurus).

### Western blotting

Cells were grown to confluence on 100 mm^2^ dishes, then lysed with RIPA buffer (Thermo) and protease inhibitor cocktail (Sigma Aldrich). Protein extracts were quantified by Pierce BCA Protein Assay kit (Thermo), loaded at 10 ug/well, run 1 hr at 150 V on a denaturing Bolt 4%–12% Bis-Tris Plus gel (Thermo) in 1X Bolt MOPS SDS buffer (Invitrogen), and transferred to a PVDF membrane (activated for 1 min in 100% methanol) (Millipore) for 1.5 hr at 10 V. After blocking for 1 hr at room temperature with 5% milk in 1X TBST (20 mM Tris base, 0.15 M NaCl, 0.1% Tween-20), membranes were probed with 1:1000 dilution of anti-MUTYH (4D10, Novus Biologicals) and as loading control, 1:1000 dilution of anti-GAPDH (D16H11, Cell Signaling Technology), followed by 1:2000 dilutions of IRDye 800CW Goat anti-Mouse and 680RD Goat anti-Rabbit IgG secondary antibodies (LI-COR). Detection was performed with Odyssey DLx Imager (LI-COR).

### Library synthesis and transduction

A wild-type *MUTYH* cDNA clone (GenBank: NM_001048174.1) was obtained from GenScript. The sequence was divided into 10 tiles; for each, a plasmid construct was synthesized and sequence-verified (Twist Bioscience) with the respective tile replaced by a 73 bp ‘stuffer’ sequence flanked by two PaqCI restriction sites (**Supplementary Table 5**). Mutant pools for each tile were designed with each codon sequentially replaced by an ‘NNN’ randomer and synthesized together as a single oPool oligo pool (IDT). Mutant libraries were PCR-amplified using tile-specific primers to add PaqCI sites, then cloned in place of the stuffer sequence in the corresponding plasmid using Golden Gate assembly with PaqCI and T4 DNA ligase (NEB), and transformed into 10-beta electrocompetent *E. coli* (NEB). These were subcloned into an inducible lentiviral expression construct (pCW57.1; Addgene #41393) by conventional directional cloning with *SalI*-HF and *NheI*-HF (NEB), and transformed into Endura electrocompetent cells (Lucigen), and plasmid mini or maxipreps were prepared using ZymoPure II kits (Zymo). Lentiviral supernatants were prepared by the University of Michigan Vector Core, as previously described^56^. Knockout cells were transduced with single mutant *MUTYH* lentivirus or pooled mutant lentivirus, at low multiplicity of infection (MOI<0.1) in the presence of 8 μg/mL polybrene (Sigma Aldrich) such that each cell carries zero or one stably integrated copies of *MUTYH* cDNA. At 48 hr post-infection, transduced cells were selected with 4 ug/ml puromycin, which was maintained in all subsequent steps.

### 8-OG:A reporter synthesis

To generate a GFP-off construct, EGFP codon 35 on pcDNA3-EGFP (Addgene #13031) was mutated from GAG (encoding Glu) to TAA (stop codon) by site-directed mutagenesis. A 2258 bp fragment spanning from 176 bp upstream of the CMV promoter to 336 bp downstream of the *bgh* polyadenylation signal was then amplified by PCR. To isolate bottom strand ssDNA, forward primers included phosphorothioate linkages between the first five bases, and the resulting PCR product was treated with T7 exonuclease (NEB) to digest the protected strand^27^. The resulting ssDNA was purified on a Clean-Concentrate-5 column (Zymo) and annealed to a partial-length top strand ssDNA prepared similarly. To site-specifically incorporate 8OG on the top strand (opposite A on the bottom strand), this partial duplex was incubated with Hemo KlenTaq DNA polymerase (NEB) and 8oxo-dGTP (TriLink) as the sole deoxynucleotide. Natural dNTPs were then added (final 0.2 mM), allowing strand extension to complete an 8OG-containing linear dsDNA product, which was purified with SPRI beads prepared as described^80^. Size and purity of each component and the final reporter were determined by gel electrophoresis (0.7% agarose, 1X TBE), and 8OG incorporation was confirmed by Sanger sequencing.

### Flow cytometry

To measure 8OG repair activity in individual cell lines (parental WT, KO, or single *MUTYH* variants), 24-well plates were seeded at 10^5^ cells/cm^2^. Variant *MUTYH* expression was induced with 10 pg/ml doxycycline (Sigma Aldrich). One day after seeding, cells were co-transfected with 125 ng 8OG reporter and 125 ng mCherry plasmid (derived from p.hBACCS2-IRES-mCherry, Addgene #72892) using Lipofectamine 3000 (Thermo). At 72h post-transfection, cells were trypsinized, washed, and stained for viability with DAPI. Out of viable transfected cells (DAPI-, mCherry+), repair-positive (GFP+) and negative (GFP-) percentages were quantified with a ZE5 Cell Analyzer flow cytometer (BioRad). The threshold beyond which cells were deemed repair-positive was set at the 95^th^ percentile of the GFP signal in *MUTYH* KO cells transfected with only the mCherry construct. Mutant libraries were similarly transfected (scaled to 10 cm dishes, each using 1.8ug each of 8OG reporter and mCherry plasmid), with three replicates per tile. Repair-positive and negative fractions were bulk-sorted with a Bigfoot Spectral Cell Sorter (Thermo), and the resulting cells were pelleted and stored at -20ºC for genomic DNA isolation. For each replicate, a fraction of 6×10^6^ mutant library-containing cells was prepared in parallel and without treatment as a pre-sort sample.

### Mutant library sequencing

Genomic DNA from pre-sort and sorted cell populations was isolated using CleanNA Blood and Tissue magnetic bead kit (Bulldog Bio). To amplify integrated *MUTYH* mutant libraries, ‘outer PCR’ was performed with primers flanking *MUTYH* cDNA using PrimeSTAR GXL kit (Takara). Each PCR reaction contained up to 500 ng gDNA as template; multiple reactions were performed and pooled as needed to consume the full amount of gDNA purified from each sorted sample. Outer PCR products were tagmented^81^ to produce random fragment libraries, and shallowly sequenced to confirm an absence of large-scale rearrangements and unintended mutations.

To quantify mutation frequency in sorted cells, mutant libraries were subjected to deep amplicon tile sequencing^82^ as described^56^. Briefly, 1ul of outer PCR library product was diluted 1:25 and amplified in a limited (≤10) cycle-PCR, with primers directed at constant bases flanking the targeted tile within *MUTYH* cDNA. Primer 5’ tails contained constant sequencing adaptor stub sequences. Each resulting product was then diluted and further amplified to add sequencing adaptors and sample-specific unique dual 10bp index sequences. Amplicon size was verified by electrophoresis on 6% TBE native PAGE gel (Thermo) stained with SYBR Gold (Thermo). Indexed libraries were pooled, purified with SPRI beads, and sequenced with paired-end 150 bp reads on NovaSeqX (Illumina) and/or Singular G4 (Singular Genomics) sequencers.

### Function score calculation

Paired end reads were overlapped and an error-corrected consensus was taken by PEAR^83^, then aligned to a *MUTYH* cDNA reference with bwa mem^84^. Variant counts were tabulated by a pipeline implemented in Snakemake (https://github.com/kitzmanlab/tileseq/). Amino acid level counts were processed by Rosace^32^ to compute a per-mutant ‘fitness’ score and a local false sign rate (lfsr) score akin to a false discovery rate^85^. To normalize between tiles, each variant score *s*_*i*_ was transformed into a log-likelihood ratio *r*_*i*_ for its membership in synonymous versus nonsense variants’ score distributions: 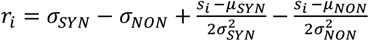, where the *µ*_*SYN*_, *σ*_*SYN*_ are the fitness score mean and standard deviation of the synonymous variants in that tile, and *µ*_*NON*_, *σ*_*NON*_ are the same for nonsense variants (codons 1-471 only). The resulting values were further shifted and scaled so to center synonymous scores at 0, and nonsense scores at -1: *a*_*i*_ = *r*_*i*_/(med_syn − med_non), where med_syn and med_non are per-tile medians of *r*_*i*_ for synonymous and nonsense variants (the latter only within codons 1-471), respectively.

### Variant classification

We classified variants as loss-of-function (LOF) if their scores fell below the 97.5^th^ percentile of nonsense variants (score≤-0.593) and had lfsr<0.05, while neutral (NEU) variants were those with scores above the 2.5^th^ percentile of synonymous variants (score≥-0.237) or which had lfsr≥0.05. Variants were considered intermediate (INT) if their scores fell between the two score thresholds (−0.593, -0.237) and had lfsr<0.05. Variants with a starting frequency below 300 counts per million (n=347, 3.5% of variants) were marked as low-abundance and removed. Splicing effects of exonic SNVs were predicted by SpliceAI^33^, and variants with deltaMax score ≥ 0.3 were considered to be splicing-disruptive.

### Clinical variants

Clinical variants and interpretations were downloaded from ClinVar on December 8, 2024 and filtered to exonic SNVs overlapping *MUTYH* transcript NM_001048174.1. For a stringently filtered set, at least two submissions were required per variant, and any variants with conflicting interpretations were excluded. For a more permissive set, conflicting interpretations were allowed only if at least two of the submitted interpretations were non-VUS, and if the non-VUS interpretations were in the same direction (e.g., benign + likely benign would be allowed; likely pathogenic + likely benign would not). For the purposes of evaluating classification accuracy, likely pathogenic and pathogenic calls were grouped together as positives, and likely benign and benign variants were grouped together as negatives.

Clinical and demographic information was obtained for patients found to carry *MUTYH* variants during cancer predisposition multi-gene panel testing at Ambry Genetics. To examine association with MAP phenotype, individuals were selected with either known pathogenic variant (founder mutations Y151C or G368D, or a truncating mutation) and a missense VUS, or with two known pathogenic variants. Several criteria were applied to select patients with phenotypes consistent with MAP: onset of colorectal cancer (excluding microsatellite unstable tumors) before age 60 years, or with polyp count ≥10 at any age. Individuals of age ≥ 50 years and with fewer than 10 polyps, or no documented history of polyps, were taken as negative for MAP. Other individuals (principally those age of age ≤ 60 years and with no history of polyps or CRC) were taken as phenotypically uninformative. Association with MAP risk was tested by Cox proportional hazards regression (R package survival) and plotted with the survminer package in R.

### Protein sequence and structure analysis

AlphaFold3^38^ was used to predict the structure of human MUTYH, taking as input the 521-amino acid protein sequence (NP_001041639), along with complementary DNA strands with an 8OG:A mismatch (5’-TGAGAC/8OG/GGG-3’ and 5’-AGTCCCAGTCTCA-3’), and a Zn2+ ion. Protein structure visualization and analysis were performed with UCSF ChimeraX (version 1.9)^86^. The matchmaker command was used to align the predicted structure to a crystal structure of mouse MUTYH (pdb ID 7ef8)^39^; after alignment, the [4Fe-4S] cluster was copied from the latter, as it is not an available ligand in the AlphaFold3 server. Secondary structures were computed by dssp^87^ and surface accessibility was computed using the ‘measure sasa’ command, each as implemented in ChimeraX. FoldX (version 5.1)^40^ was used to predict changes in folding free energy of all missense mutations in MUTYH.

GEMME scores^88^ were computed taking as input a multiple sequence alignment obtained by the ConservFold pipeline^89^ and ColabFold^90^. The per-residue mean GEMME score was taken^91^ as a measure of evolutionary conservation. Using python statsmodels^92^ version 0.14.4, a logistic regression model was fitted with the per-residue fraction of missense variants scoring as INT or LOF, with features including the relative sasa score and per-residue mean scores from: FoldX, AlphaMissense^93^, ESM1v^94^, and GEMME.

## Data availability

All single-codon and single-base function scores and associated annotations are supplied in Supplementary Tables 1 and 2). Raw sequencing data and processed variant scores are available at NCBI GEO (accession GSE290379).

## Acknowledgements

We thank Bala Burugula, Emily Kirk, and Sajini Jakakody for sequencing and cloning assistance; Sebastian Vishnopolska and Anthony Scott for helpful feedback; Cynthia Martinez, Cassidy Carraway and Taylor Coleman for assistance with clinical data retrieval and curation. This work was supported by the National Institute of General Medical Sciences (R35GM153286 to J.O.K) and an American Cancer Society Postdoctoral Fellowship (to S.L.H.).

## Supplementary Materials

Supplementary Figures 1-9

Supplementary Tables 1-5

