## Supplementary Figures for "Saturation mapping of *MUTYH* variant effects using DNA repair reporters"

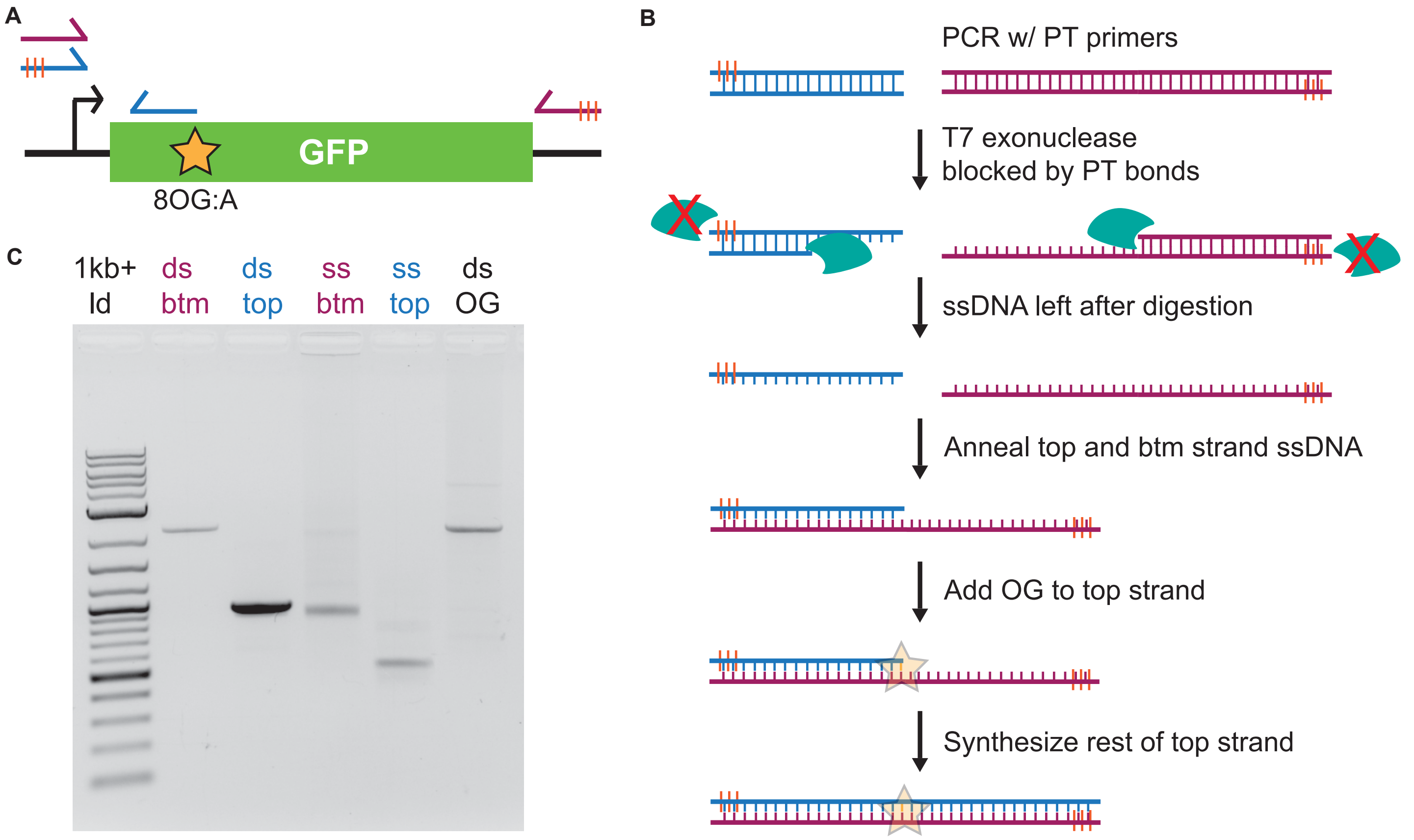


**Supplemental Figure 1: 8OG reporter generation. (A)** Schematic of GFP 8OG reporter. Star indicates 8OG on top strand at codon 35, opposite an A. Blue primers amplify up to codon 35. Red primers amplify whole construct. Orange bars on the primers indicate phosphorothioate (PT) bonds between the first five 5' nucleotides which render the resulting strands exonuclease-resistant. **(B)** Overview of reporter generation. PCR amplification with PT-containing primers produces amplicons with PT bonds at the 5' end of top or bottom strand, depending on primer pair. Incubation of each PCR amplicon with T7 exonuclease results in digestion of the non-PT-containing strand. Top and bottom strands are annealed, producing a partial duplex, with the top strand 3' most base positioned immediately before codon 35 (ATT on the ssDNA bottom strand). 8OG incorporation occurs by strand extension with deoxy-8OGTP as the only nucleotide. Natural dNTPs are then spiked into the reaction, allowing DNA polymerase to synthesize the rest of the top strand. **(C)** Agarose gel of PCR products, T7 exonuclease digestion products, and assembled reporter.

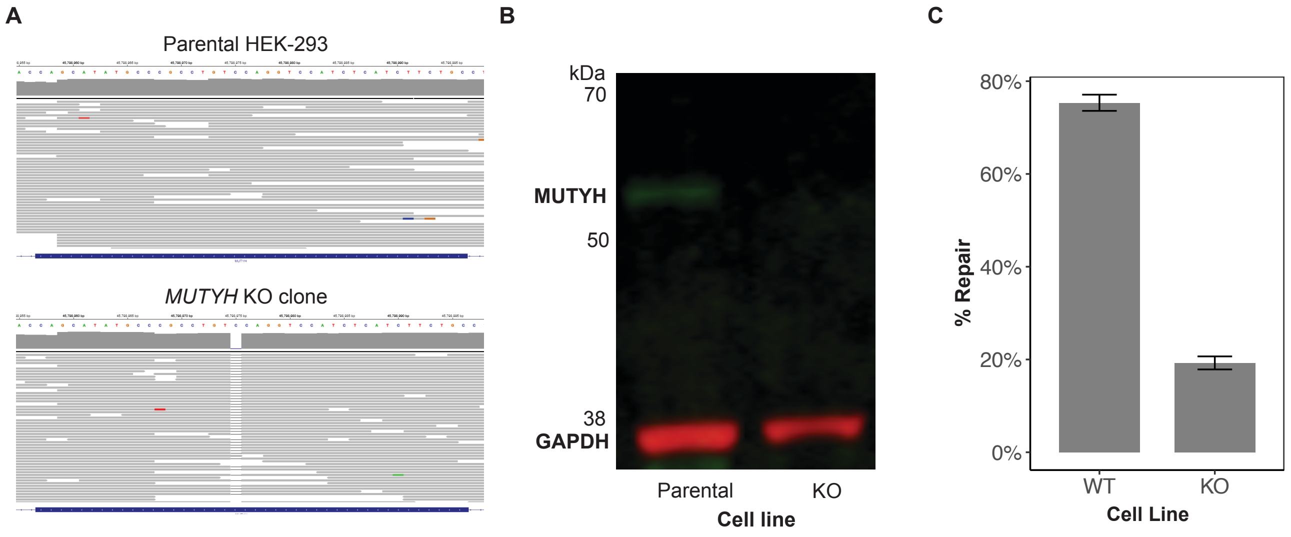


**Supplemental Figure 2: *MUTYH* KO cell line validation. (A)** Deep sequencing to genotype *MUTYH* Cas9 target site in parental (upper) and *MUTYH* KO (lower) HEK-293 cells. Editing created a homozygous 1-bp deletion (chr1:45798975, hg38 coordinates) resulting in frameshift and premature truncation (c.290delC, p.Arg97fsSer20*). **(B)** Western blot confirms loss of MUTYH expression in KO cells. **(C)** Quantification of 8OG:A repair of HEK293 WT and KO cells. % Repair is calculated as total GFP+ cells divided by total mCherry+ cells.

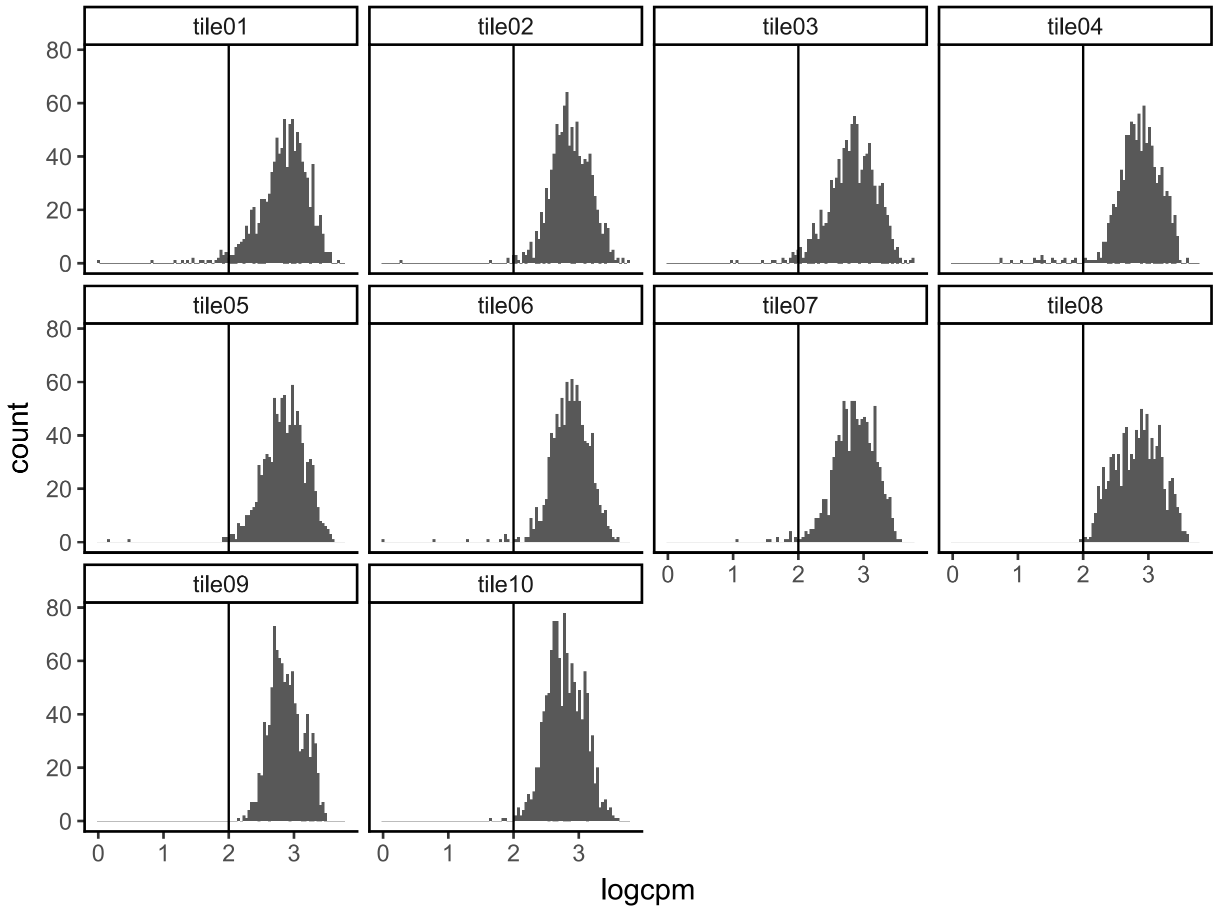


**Supplementary Figure 3. Mutant library uniformity.** Histograms of *MUTYH* variant abundance, $\log_{10}$(counts/million counts), within plasmid libraries for each mutagenesis tile. Cutoff line is at 100 cpm (1/10,000).

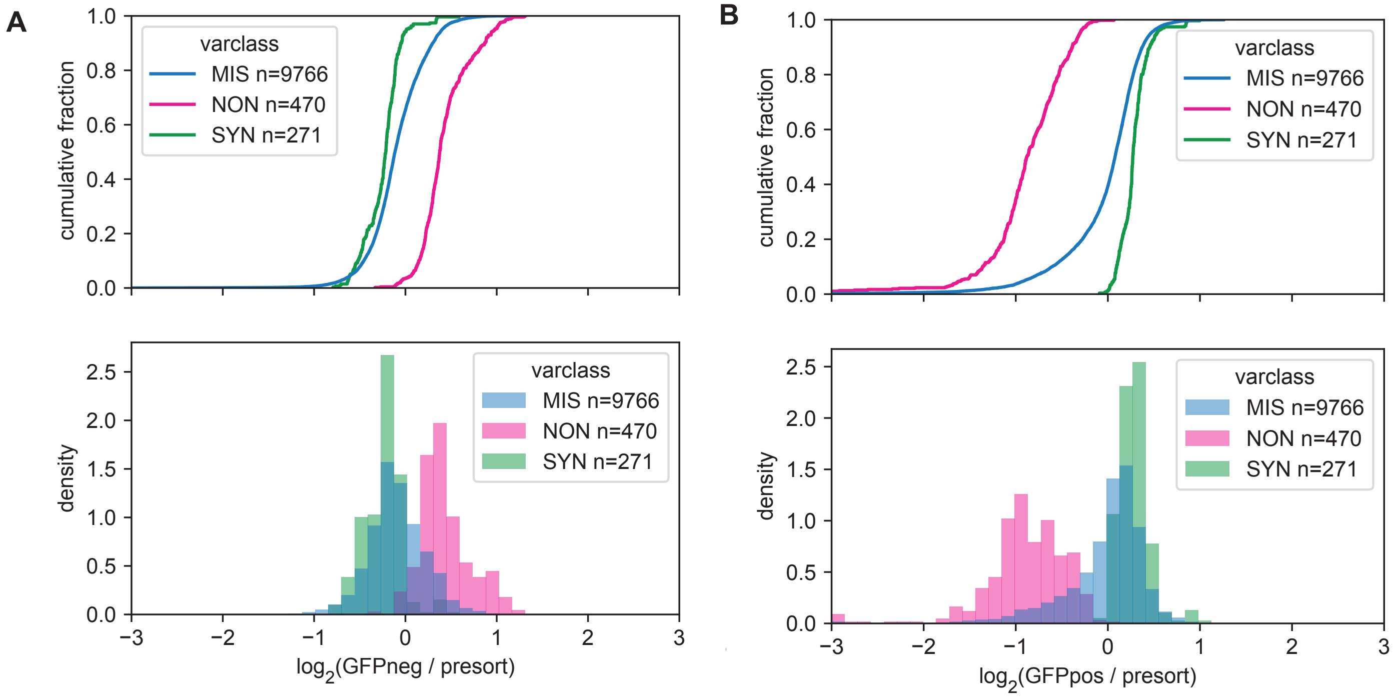


**Supplementary Figure 4. Variant enrichment and depletion.** Empirical cumulative distributions (above) and histograms (below) of variant function score, for **A.** GFP-negative sorts (in which nonsense variants are enriched), versus the pre-sorted population, and **B.** GFP-positive sorts (in which nonsense variants are depleted), versus the pre-sorted population. Distributions are shaded by functional category; nonsense variants at codon ≥ 472 are excluded.

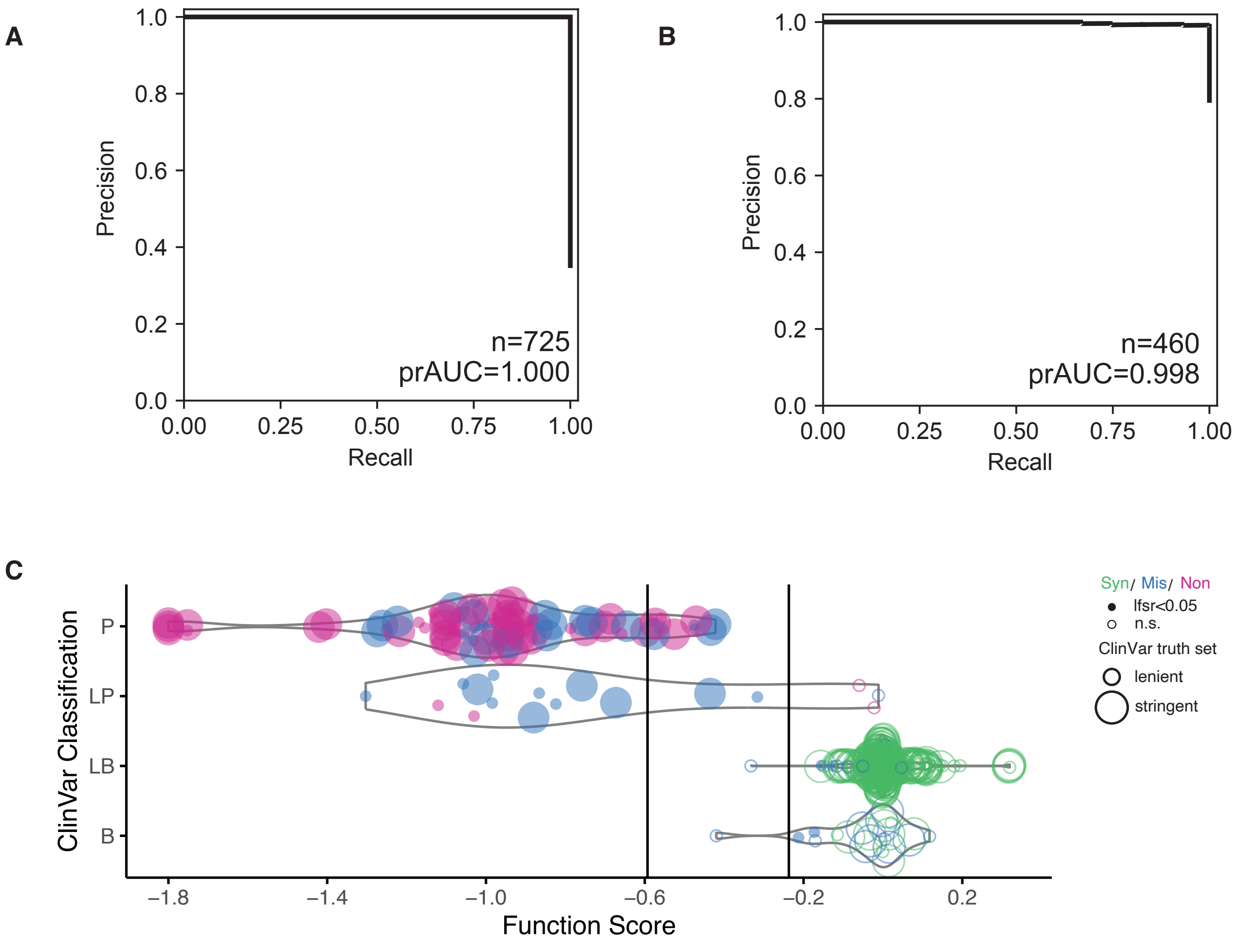


**Supplementary Figure 5. Classification performance using function scores.** **(A)** Precision-recall curve showing perfect separation of 725 nonsense and synonymous variants in codons 1-471. **(B)** Precision-recall curve showing near-perfect classification performance on truth set ClinVar *MUTYH* SNVs with lenient filtering (resoling conflicting entries as pathogenic or benign when there were ≥2 non-VUS submission records). **(C)** Function scores plotted for lenient filtered ClinVar set, plotted and shaded as in **Fig. 4A**. Point size whether each variant passed stringent filters or not.

**A**

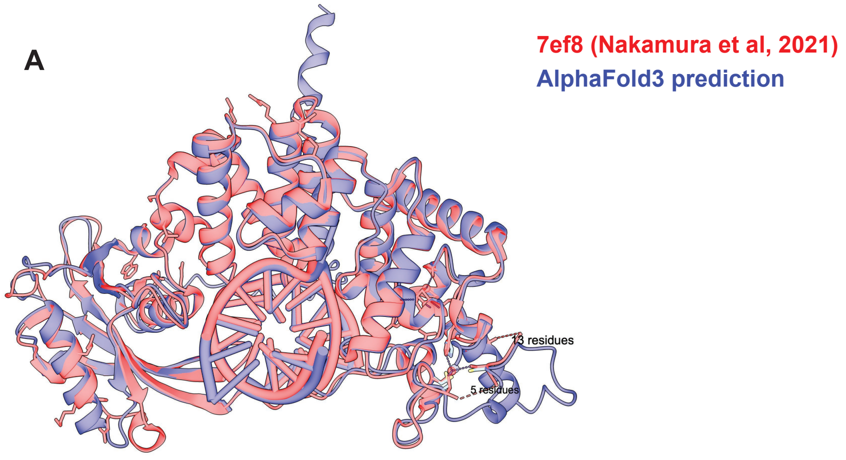


**B**

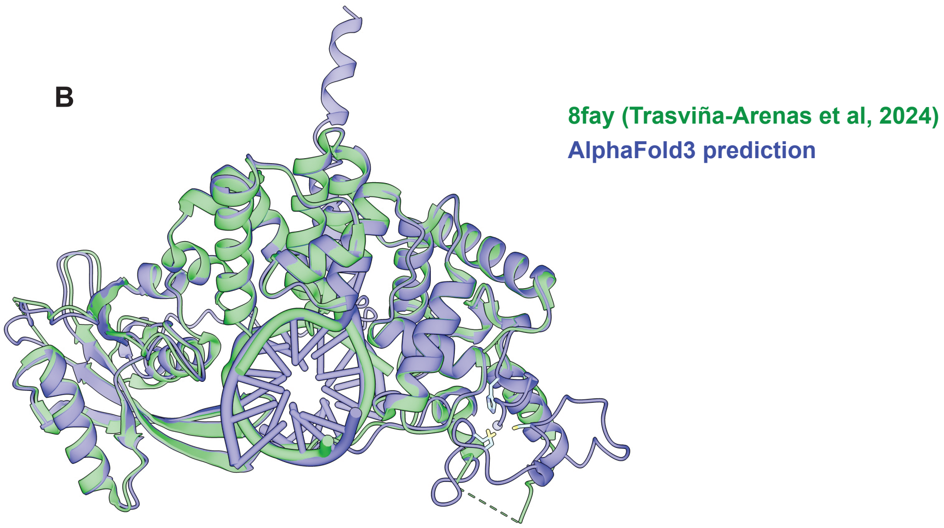


**Supplementary Figure 6. Comparison of predicted and experimentally resolved MUTYH structures.** Structural alignments from ChimeraX matchmaker of AlphaFold3-predicted structure of 521-amino acid human MUTYH sequence in complex with 8OG:A bearing DNA, along with **(A)** mouse MUTYH (7ef8; rmsd=0.73A), and **(B)** human MUTYH (8fay; rmsd=0.70A).

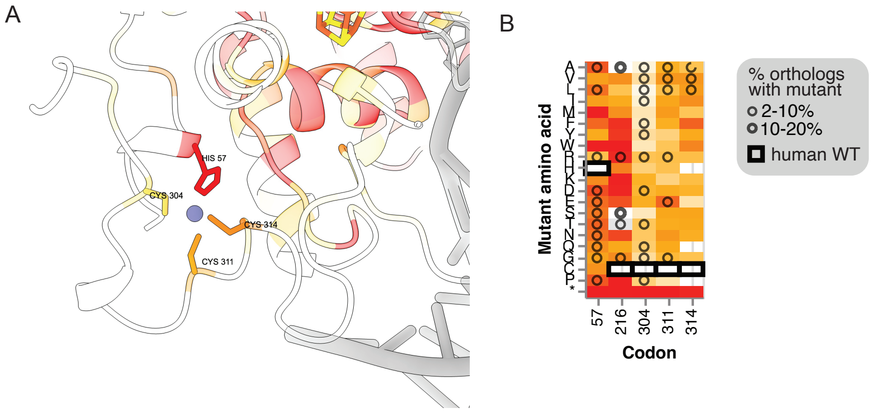


**Supplementary Figure 7. Constraint at $Zn^{2+}$ binding motif. (A)** Zinc-binding motif within AlphaFold3-predicted structure of human MUTYH, with residues shaded by average missense constraint score, as in Figure 5. **B.** Heatmap detail with function score at previously proposed zinc-binding residues, shaded as in Figure 3. Dark rectangles denote wild type residues, and circles indicate residues in aligned MUTYH orthologs.

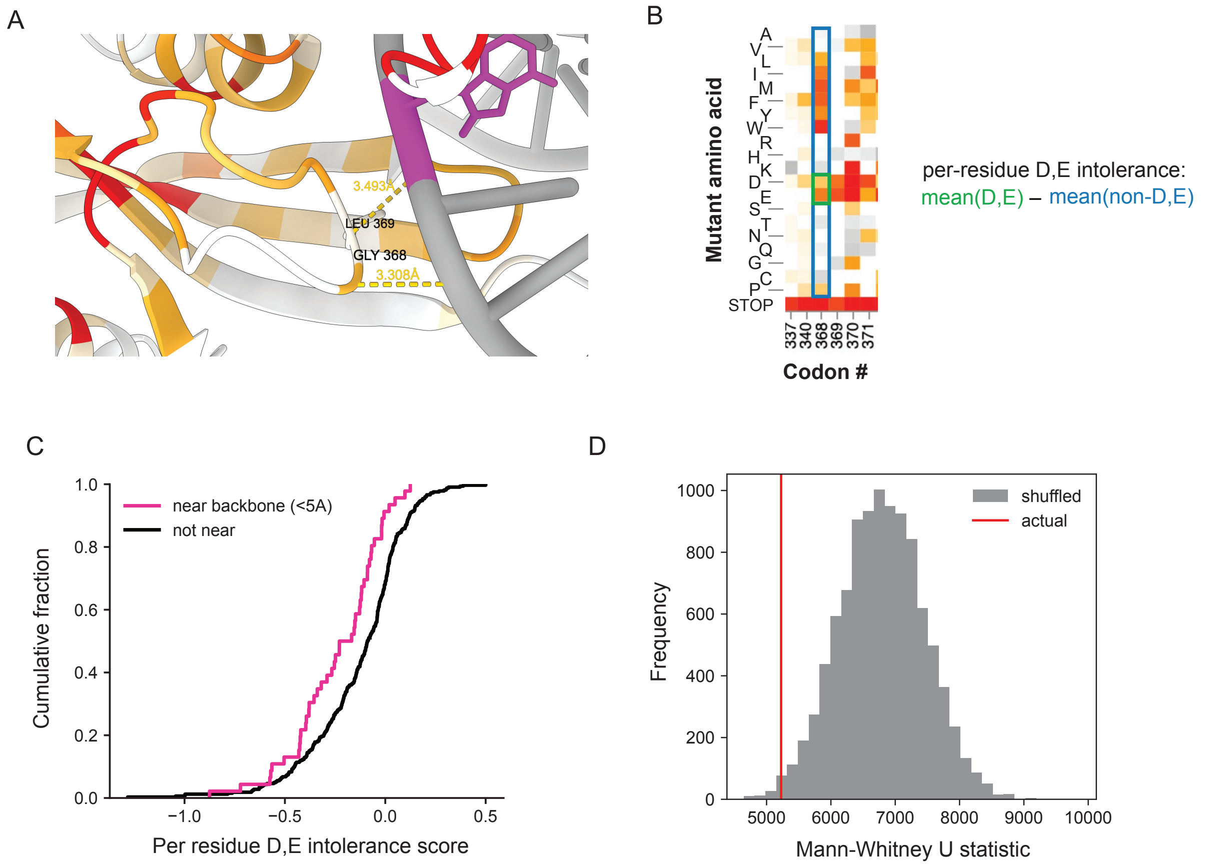


**Supplementary Figure 8. Intolerance to negatively charged amino acids near DNA backbone. (A)** View of residues G368 and L369, with distances to DNA backbone marked (<3.5Å for each). **(B)** Heatmap illustrating per-residue D,E intolerance score, the difference in mean function scores of D or E mutations and non-D/E mutations. **(C)** Cumulative distribution of D,E intolerance scores for residues within 5Å of the DNA backbone (pink) and all other residues within codons 57-472. **(D)** Mann-Whitney U test statistics resulting from comparison of D,E intolerance scores among backbone-proximal residues vs distal residues randomly sampled without replacement with similar surface exposure to the backbone-proximal set; actual test (red) is overlaid on a histogram of 10,000 random samples (gray).

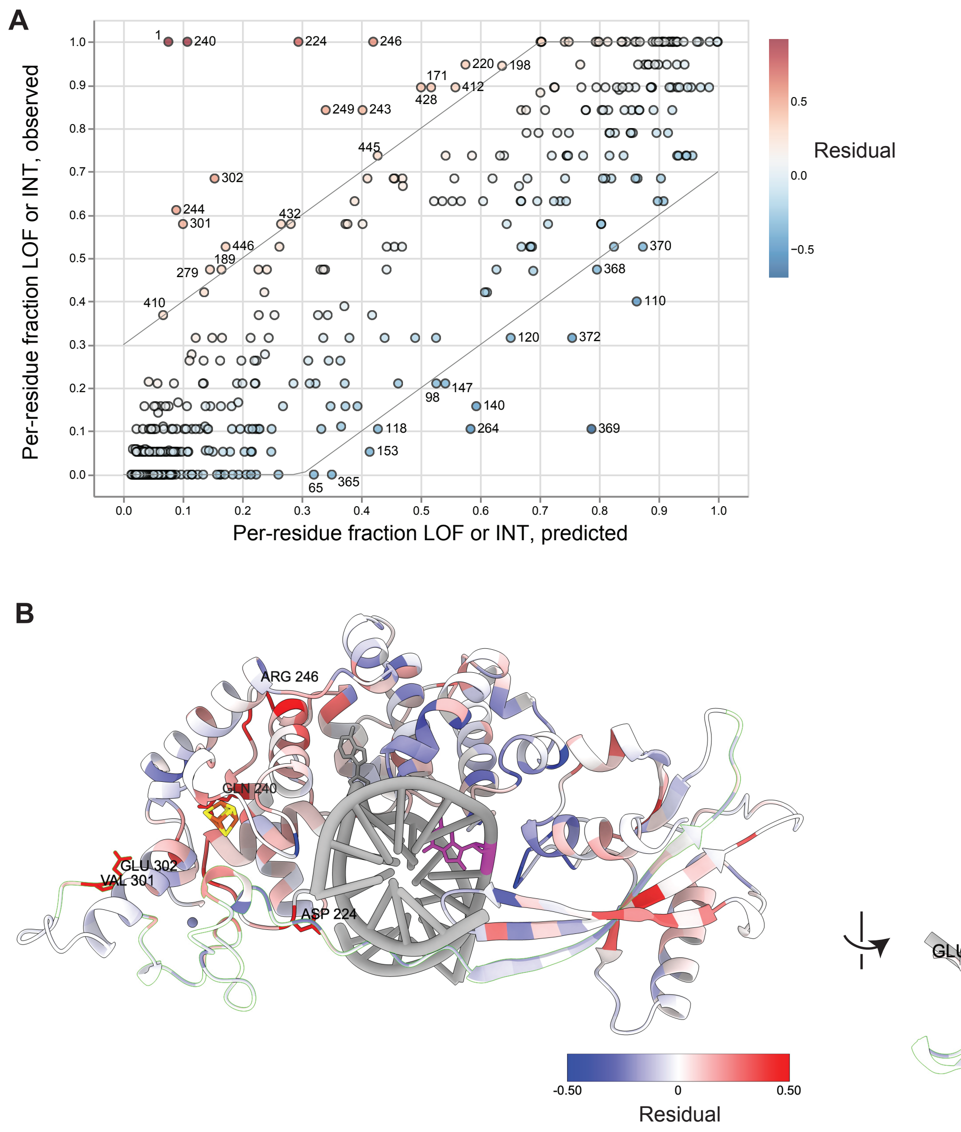


**Supplementary Figure 9. Constraint at residues not predicted by evolutionary/structural features. (A)** Per-residue constraint (fraction of missense mutations scoring as LOF or INT) plotted as observed (y) versus that predicted from logistic regression model fitted with structural/evolutionary features (SASA, FoldX, AlphaMissense, GEMME, and ESM-1 scores). Codon position numbers are shown for residues with absolute value of residual ≥ 0.30. Points are shaded by the signed residual value; blue: predicted to be more constrained than observed; red: observed constraint greater than predicted. **(B)** AlphaFold3 structure of MUTYH shaded by per-residue residual score, shaded as in (A). Several residues with low-predicted but high-observed constraint are labeled (D224, Q240, R246, V301, E302); the interdomain connector loop is indicated with a light-green border.
